# Sensory experience modulates the reorganisation of temporal auditory regions for executive processing

**DOI:** 10.1101/2021.02.08.430248

**Authors:** Barbara Manini, Valeria Vinogradova, Bencie Woll, Donnie Cameron, Martin Eimer, Velia Cardin

## Abstract

Crossmodal plasticity refers to the reorganisation of sensory cortices in the absence of their main sensory input. Understanding this phenomenon provides insights into brain function and its potential for change and enhancement. Using fMRI, we investigated how early deafness influences crossmodal plasticity and the organisation of executive functions in the adult human brain. Results from a range of visual executive function tasks (working memory, task switching, planning, inhibition) show that deaf individuals specifically recruit “auditory” regions during task switching. Neural activity in superior temporal regions, most significantly in the right hemisphere, are good predictors of behavioural performance during task switching in the deaf group, highlighting the functional relevance of the observed cortical reorganisation. Our results show executive processing in typically sensory regions, suggesting that the development and ultimate role of brain regions are influenced by perceptual environmental experience.

## Introduction

Sensory systems feed and interact with all aspects of cognition. As such, it is likely that developmental sensory experience will impact the organisation of higher-order cognitive processes such as executive functions. Here we studied executive processing in early deaf individuals in order to understand the influence of early sensory experience on higher-order cognition and neural reorganisation.

Executive functions are higher-order cognitive processes responsible for flexible and goal-directed behaviours, which have been associated with activity in frontoparietal areas of the brain.^1^ However, studies on deafness have shown reorganisation for visual working memory in regions typically considered to be part of the auditory cortex.^2–4^ These working memory responses in auditory regions suggest that, in the absence of early sensory stimulation, a sensory region can change its function as well as the perceptual modality to which it responds.^5,6^ The adaptation of sensory brain regions to processing information from a different sensory modality is known as crossmodal plasticity.^6–18^ The presence of working memory responses in deaf auditory regions takes the concept of crossmodal plasticity further, suggesting that, in the absence of early auditory stimulation, there is a shift from sensory to cognitive processing in deaf auditory regions. If this is the case, it would suggest that cortical functional specialisation for sensory or cognitive processing is partially driven by environmental sensory experience. The aim of our study is to elucidate the role of the deaf auditory cortex in executive functions, in order to understand how sensory experience impacts cognitive processing in the brain. Specifically, we tested whether deaf auditory regions are involved in cognitive control or whether they have a role in specific subcomponents of executive functions.

To address our aims, we conducted an fMRI experiment in deaf and hearing individuals. Participants performed tasks tapping into different executive functions: working memory, planning, switching, and inhibition. If the deaf auditory cortex has a role in cognitive control, we would expect all tasks to recruit this region. However, if deaf auditory areas are involved in specific subcomponents of executive functioning, these regions will be differentially activated by each of the tasks. If neural activity in the reorganised auditory cortex can predict behavioural performance in deaf individuals, this will corroborate the functional significance of such plasticity effect.^19,20^

## Materials and Methods

### Participants

There were two groups of participants (see summary demographics in Supplementary Table 1-2):

a. 29 congenitally or early (before 3 years of age) severely-to-profoundly deaf individuals whose first language is British Sign Language (BSL) and/or English (Supplementary Table 3). We recruited a larger number of deaf participants to reflect the language variability of the deaf population in the UK, as discussed in the “Language assessment” section. Datasets from three deaf participants were excluded from all analyses due to excessive motion in the scanner. One participant was excluded because they only had a mild hearing loss in their best ear (pure-tone average (PTA) less than 25dB). In total, 25 deaf participants were included in the analysis of at least one executive function task (see Supplementary Table 4 for details on exclusion).
b. 20 hearing individuals who are native speakers of English with no knowledge of any sign language.

Deaf and hearing participants were matched on age, gender, nonverbal intelligence, and visuospatial working memory span (Supplementary Table 2).

All participants gave written informed consent. All procedures followed the standards set by the Declaration of Helsinki and were approved by the ethics committee of the School of Psychology at the University of East Anglia (UEA) and the Norfolk and Norwich University Hospital (NNUH) Research and Development department.

Participants were recruited through public events, social media, and participant databases of the UCL (University College London) Deafness, Cognition and Language Research Centre (DCAL) and the UEA School of Psychology. Participants were all right-handed (self-reported), had full or corrected vision, and no history of neurological conditions. All participants were compensated for their time, travel, and accommodation expenses.

### General procedure

Participants took part in one behavioural and one scanning session. The sessions took place on the same or different days.

The behavioural session included:

a. *Standardised nonverbal IQ and working memory tests:* the Block Design subtest of the Wechsler Abbreviated Scale of Intelligence^21^ (WASI) and the Corsi Block-tapping test^22^ implemented in PEBL software^23^ (http://pebl.sourceforge.net/).
b. *Language tasks:* four tasks were administered to assess language proficiency in English and BSL in deaf participants (see the “Language assessment” section below).
c. *Pre-scanning training:* the training session ensured that participants understood the tasks and reached accuracy of at least 75%. The tasks were explained in the participant’s preferred language (English or BSL).
d. *Audiogram screening*: pure-tone averages (PTAs) were used to measure the degree of deafness in deaf participants. Copies of audiograms were provided by the participants from their audiology clinics or were collected at the time of testing using a Resonance R17 screening portable audiometer. Participants included in the study had a mean PTA greater than 75dB averaged across the speech frequency range (0.5, 1, 2kHz) in both ears (mean=93.66±7.79dB; range: 78.33-102.5dB). Four participants did not provide their audiograms but they were all congenitally severely or profoundly deaf and communicated with the researchers using BSL or relying on lipreading.

During the scanning session, fMRI data were acquired while participants performed four visual executive function tasks on working memory, planning, switching, and inhibition (see details below). The order of the tasks was counterbalanced across participants.

### Experimental design

All tasks were designed so that each had one condition with higher executive demands (Higher Executive Function; HEF) and one with lower demands (Lower Executive Function; LEF) (Figure 1).

**Figure 1.**
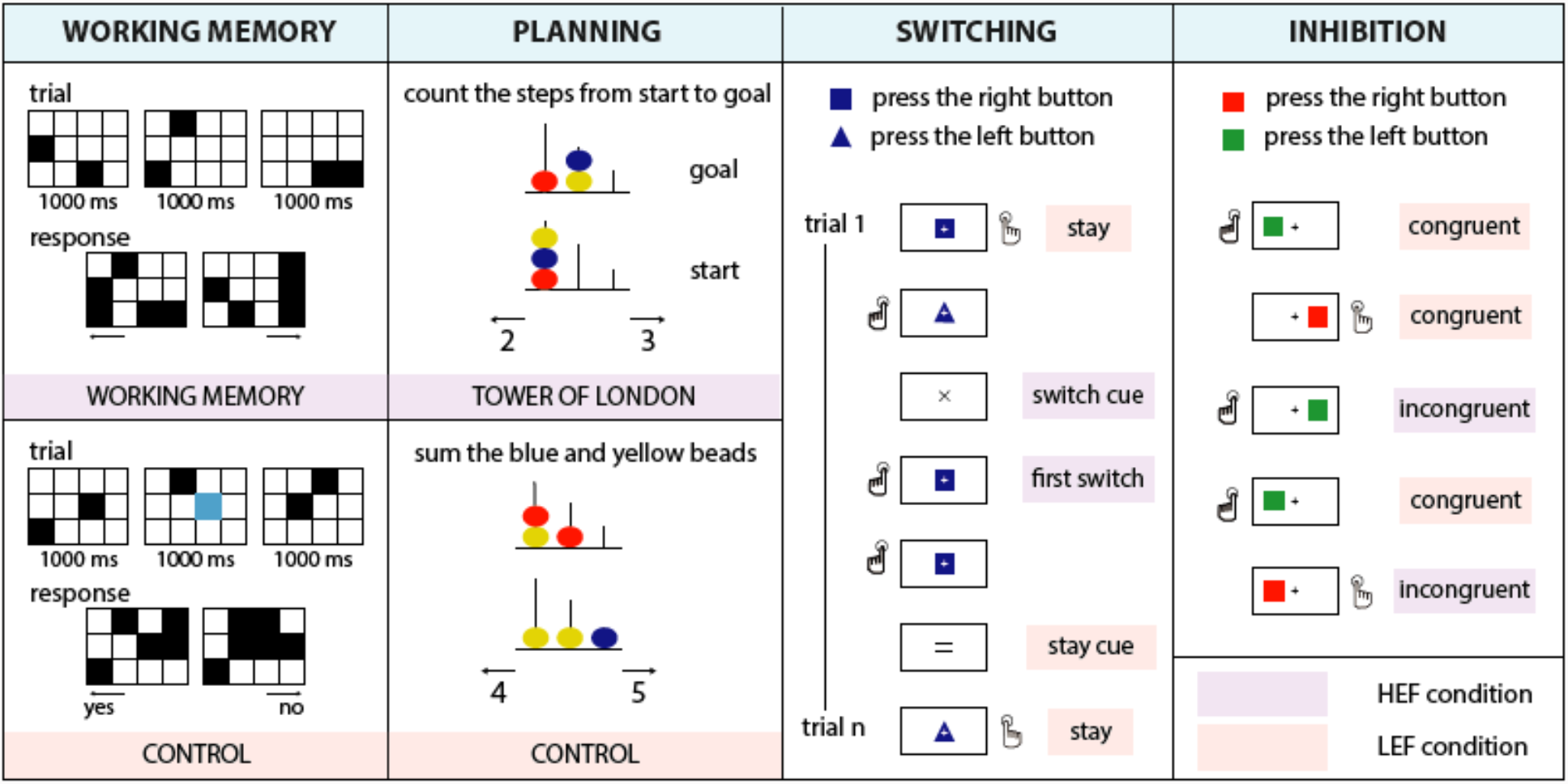
Executive function tasks. Each task had a higher executive demands condition (HEF=Higher Executive Function, purple) and a lower executive demands condition (LEF=Lower Executive Function, peach). See Methods for details of the design.

#### Working memory

We used a visuospatial working memory task^24,25^ (Figure 1) contrasted with a perceptual control task. A visual cue (1500ms) indicated which task participants should perform. The cue was followed by a 3×4 grid. Black squares were displayed two at a time at random locations on the grid, three times, for a total of 1000ms. In the HEF condition, participants were asked to memorise the six locations. Then they indicated their cumulative memory for these locations by choosing between two grids in a two-alternative, forced-choice paradigm via a button press. The response grids were displayed until the participant responded or for a maximum of 3750ms. In the control condition (LEF), participants indicated whether a blue square was present in any of the grids, ignoring the configuration of the highlighted squares. Trials were separated by an inter-trial interval (ITI) with duration jittered between 2000-3500ms. Each experimental run had 30 working memory trials and 30 control trials.

#### Planning

We used a computer version of the classic Tower of London task^26,27^ (Figure 1). In each trial, two configurations of coloured beads placed on three vertical rods appeared on a grey screen, with the tallest rod containing up to three beads, the middle rod containing up to two beads, and the shortest rod containing up to one bead. In the Tower of London condition (HEF), participants had to determine the minimum number of moves needed to transform the starting configuration into the goal configuration following two rules: 1) only one bead can be moved at a time; 2) a bead cannot be moved when another bead is on top. There were four levels of complexity, depending on the number of moves required (2, 3, 4, and 5). In the control condition (LEF), participants were asked to count the number of yellow and blue beads in both displays. For both conditions, two numbers were displayed at the bottom of the screen: one was the correct response and the other was incorrect by +1 or -1. Participants answered with their left hand when they chose the number on the left side of the screen, and with their right hand when their choice was on the right. The maximum display time for each stimulus was 30 seconds. The duration of the ITI was jittered between 2000-3500ms. There were 30 trials in the Tower of London condition and 30 trials in the control condition.

#### Switching

In this task, participants had to respond to the shape of geometric objects, i.e., a rectangle and a triangle^28,29^ (Figure 1). At the beginning of the run, participants were instructed to press a key with their left hand when they saw a rectangle and with their right hand when they saw a triangle. Each block started with a cue indicating that the task was to either keep the rule they used in the previous block (“stay” trials; LEF) or to switch it (“switch” trials; HEF). In the switch trials, participants had to apply the opposite mapping between the shape and the response hand. Each block included the presentation of the instruction cue (200ms), a fixation cross (500ms), and two to five task trials. During each trial, a geometrical shape (either a blue rectangle or a blue triangle) was shown at the centre of the screen until the participant responded for a max of 1500ms. Visual feedback (500ms) followed the participant’s response. There were 230 trials in 80 blocks of either the LEF (40) or HEF (40) condition. The analysis for the HEF condition only included the first trial of the switch block (see below).

#### Inhibition

To study inhibitory control, we used Kelly and Milham’s version of the classic Simon task (https://exhibits.stanford.edu/data/catalog/zs514nn4996).^30^ A square appeared on the left or the right side of the fixation cross. The colour of the squares was the relevant aspect of the stimuli, with their position irrelevant for the task. Participants were instructed to respond to the red square with the left hand and the green square with the right hand. In the congruent condition (LEF), the button press response was spatially congruent with the location of the stimuli (e.g. the right-hand response for a square appearing on the right side of the screen) (Figure 1). In the incongruent condition (HEF), the correct answer was in the opposite location in respect to the stimulus. Half of the trials were congruent, and half were incongruent.

Each stimulus was displayed for 700ms, with a response window of up to 1500ms. The ITI was 2500ms for most trials, with additional blank intervals of 7.5 seconds (20), 12.5 seconds (2), and 30 seconds (1). Participants completed 1 or 2 runs of this task, each consisting of a maximum of 200 trials.

### Statistical analysis of behavioural performance in executive function tasks

Averaged accuracy (%correct) and reaction time (RT) were calculated. For each participants’ set of RTs, we excluded outlier values that were more than 1.5 interquartile ranges below the first quartile or above the third quartile of the data series. Differences between groups on accuracy or RT were investigated with repeated-measures ANOVAs with between-subjects factor group (hearing, deaf) and within-subjects factor condition (LEF, HEF).

In the switching task, the accuracy switch cost (SwitchCost_ACC_) was calculated as the difference in the percent of errors (%errors) between the first switch trial of a switch block and all stay trials. RT switch cost (SwitchCost_RT_) was calculated as the difference in RT between the first switch trial of a switch block and all stay trials.

In the inhibition task, the Simon effect was calculated as the difference in %errors or RT between the incongruent and congruent trials.

### Image acquisition

Images were acquired at the Norfolk and Norwich University Hospital (NNUH) in Norwich, UK, using a 3 Tesla wide bore GE 750W MRI scanner and a 64-channel head coil. Communication with the deaf participants occurred in BSL through a close-circuit camera, or through written English through the screen. An intercom was used for communication with hearing participants. All volunteers were given ear protectors. Stimuli were presented with PsychoPy software^31^ (https://psychopy.org) through a laptop (MacBook Pro, Retina, 15-inch, Mid 2015). All stimuli were projected by an AVOTEC’s Silent Vision projector (https://www.avotecinc.com/high-resolution-projector) onto a screen located at the back of the magnet’s bore. Participants watched the screen through a mirror mounted on the head coil. Button responses were recorded via fORP (Fiber Optic Response Pads) button boxes (https://www.crsltd.com/tools-for-functional-imaging/mr-safe-response-devices/forp/). Functional imaging data were acquired using a gradient-recalled echo (GRE) EPI sequence (50 slices, TR=3,000ms, TE=50ms, FOV=192×192mm, 2mm slice thickness, distance factor 50%) with an in-plane resolution of 3×3mm. The protocol included six functional scans: 1 resting state scan (reported in a different manuscript) and five task-based fMRI scans (working memory: 11 minutes, 220 volumes; planning: 11.5 minutes, 230 volumes; switching: 10.5 minutes, 210 volumes; inhibition: two runs of 10 minutes, 200 volumes each). Some participants did not complete all functional scans (Supplementary Table 4). An anatomical T1-weighted scan (IR-FSPGR, TI=400ms, 1mm slice thickness) with an in-plane resolution of 1×1mm was acquired during the session.

Raw B0 field map data were acquired using a 2D multi-echo GRE sequence with the following parameters: TR=700ms, TE=4.4 and 6.9ms, flip angle=50°, matrix size=128×128, FOV=240mm×240mm, number of slices=59, thickness=2.5mm, and gap=2.5mm. Real and imaginary images were reconstructed for each TE to permit calculation of B0 field maps in Hz.^32–34^

### fMRI preprocessing

fMRI data were analysed with MATLAB 2018a (MathWorks, MA, USA) and Statistical Parametric Mapping software (SPM12; Wellcome Trust Centre for Neuroimaging, London, UK).^35^ The anatomical scans were segmented into different tissue classes: grey matter, white matter, and cerebrospinal fluid. Skull-stripped anatomical images were created by combining the segmented images using the Image Calculation function in SPM (ImCalc, http://tools.robjellis.net). The expression used was: [(i1.*(i2+i3+i4))>threshold], where i1 was the bias-corrected anatomical scan and i2, i3 and i4 were the tissue images (grey matter, white matter, and cerebrospinal fluid, respectively). The threshold was adjusted between 0.5 and 0.9 to achieve adequate brain extraction for each participant. Each participant’s skull-stripped image was normalised to the standard MNI space (Montreal Neurological Institute) and the deformation field obtained during this step was used for normalisation of the functional scans. Susceptibility distortions in the EPI images were estimated using a field map that was co-registered to the BOLD reference.^32,33^ Images were realigned using the pre-calculated phase map, co-registered, slice-time corrected, normalised, and smoothed (using an 8mm FWHM Gaussian kernel). All functional scans were checked for motion and artefacts using the ART toolbox (https://www.nitrc.org/projects/artifact_detect).

### fMRI first-level analysis

The first-level analysis was conducted by fitting a general linear model (GLM) with regressors of interest for each task (see details below). All the events were modelled as a boxcar and convolved with SPM’s canonical hemodynamic response function.

The motion parameters, derived from the realignment of the images, were added as regressors of no interest. Regressors were entered into a multiple regression analysis to generate parameter estimates for each regressor at every voxel.

#### Switching

The first trial of each switch block (HEF) and all stay trials (LEF) were modelled as regressors of interest separately for the left- and right-hand responses. The cues and the remaining switch trials were included as regressors of no interest.

#### Working memory

The conditions of interest were working memory (HEF) and control (LEF). The onset was set at the presentation of the first grid, with the duration set at 3.5 seconds (i.e., the duration of the three grids plus a 500ms blank screen before the appearance of the response screen; Figure 1). Button responses were included separately for each hand and condition as regressors of no interest.

#### Planning

Tower of London (HEF) and control (LEF) conditions were included in the model as regressors of interest, with onsets at the beginning of each trial and duration set to the trial-specific RT. Button responses were modelled separately for each hand as regressors of no interest.

#### Inhibition

Four regressors of interest were obtained by combining the visual hemifield where the stimulus appeared with the response hand (1. right visual hemifield*—*left hand; 2. left visual hemifield*—*right hand; 3. right visual hemifield*—*right hand; 4. left visual hemifield*—*left hand). Right visual hemifield*—*left hand and left visual hemifield*—*right hand were the incongruent conditions (HEF), whereas the right visual hemifield-right hand and left visual hemifield-left hand were the congruent conditions (LEF).

### Region of interest analysis

We conducted a region of interest (ROI) analysis to investigate crossmodal plasticity and differences between groups in the auditory cortex. Three auditory regions of the superior temporal cortex were included in this analysis: Heschl’s gyrus (HG), the planum temporale (PT), and the posterior superior temporal cortex (pSTC) (Figure 2). HG and the PT were defined anatomically, using FreeSurfer software^36^ (https://surger.nmr.mgh.harvard.edu). Full descriptions of these procedures can be found elsewhere^37,38^, but in short, each participant’s bias-corrected anatomical scan was parcellated and segmented, and voxels with the HG label and the PT label were exported using SPM’s ImCalc function (http://robjellis.net/tools/imcalc_documentation.pdf). Participant-specific ROIs were then normalised to the standard MNI space using the deformation field from the normalisation step of the preprocessing.

**Figure 2.**
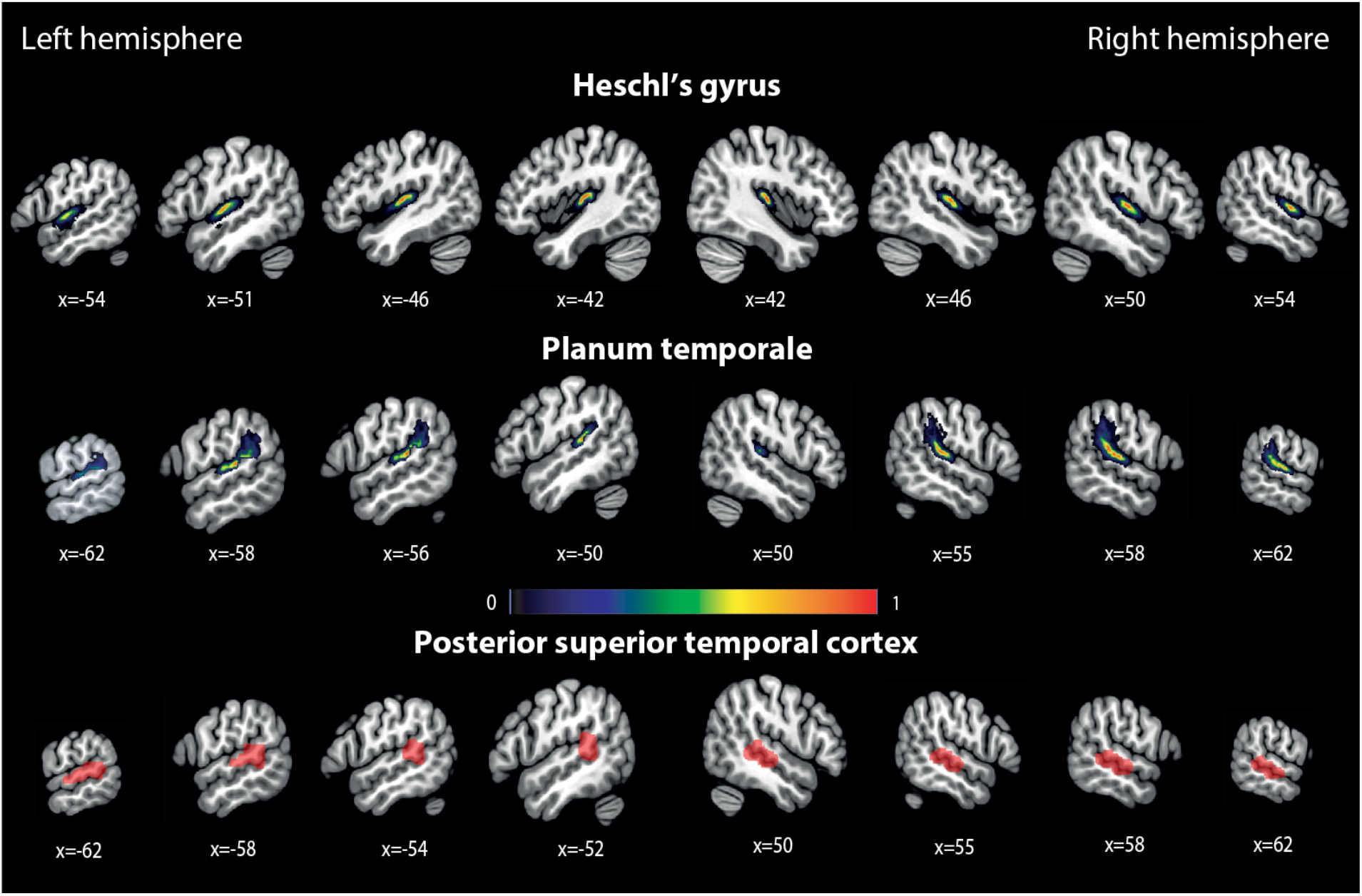
Auditory Superior Temporal ROIs A. Temporal regions included in the analysis: Heschl’s gyrus (HG), the planum temporale (PT), and the superior temporal cortex (pSTC). HG and PT were defined anatomically, in a subject-specific manner, using the FreeSurfer software package.^36^ The figure shows the overlap of all subject-specific ROIs. Common voxels between left PT and left pSTC have been subtracted from left PT (see Methods). pSTC was defined functionally, based on the findings of Cardin et al.’s study^3^ (see Methods).

pSTC was specified following findings from Cardin et al.’s study^3^, where a visual working memory crossmodal plasticity effect was found in right and left pSTC in deaf individuals [left: -59 -37 10; right: 56 -28 -1]. Right and left functional pSTC ROIs were defined using data from Cardin et al.^3^, with the contrast [deaf (working memory > control task) > hearing (working memory > control task)] (p<0.005, uncorrected).

There was an average partial overlap of 8.2 voxels (SD=6.86) between left PT and left pSTC, with no significant difference in overlap between groups (deaf: mean=9.92, SD=7.02; hearing: mean=6.05, SD=6.17). To ensure that the two ROIs were independent, common voxels were removed from left PT in a subject-specific manner. Removing the overlapping voxels did not qualitatively change the results.

### ROI analysis

Parameter estimates for each participant were extracted from each ROI using MarsBaR 0.44^39^ (http://marsbar.sourceforge.net). The data were analysed using JASP^40^ (https://jasp-stats.org), and entered into separate repeated-mixed measures ANOVAs for each task and set of ROIs. Factors in the ANOVAs on the temporal ROIs included: the between-subjects factor group (hearing, deaf) and the within-subjects factors ROI (HG, PT, pSTC), hemisphere (left, right), and condition (LEF, HEF).

The Greenhouse-Geisser correction was applied when the assumption of sphericity was violated. Significant interactions and effects of interest were explored with Student’s t-tests or Mann-Witney U-tests when the equal variance assumption was violated.

### Language assessment

We recruited a representative group of the British deaf population, who usually have different levels of proficiency in sign and spoken language. This was: 1) to study plasticity in a representative group of deaf individuals; 2) to study the relationship between language experience and the organisation of cognitive networks of the brain, which will be reported in a separate manuscript.

To assess the language proficiency of deaf participants, we chose grammaticality judgement tests measuring language skills in English and BSL. The BSL grammaticality judgement task (BSLGJT) is described in Cormier et al.’s paper^41^, and the English grammaticality judgement task (EGJT) was designed based on examples from Linebarger et al.’s paper^42^. The BSLGJT and the EGJT use a single method of assessing grammaticality judgements of different syntactic structures in English and BSL. Grammaticality judgement tests have been used in deaf participants before and have proved to be efficient in detecting differences in language proficiency among participants with varying ages of acquisition.^41,43^ Deaf participants performed both the BSL and English tests if they knew both languages, or only the English tests if they did not know BSL.

To control for potential language proficiency effects, we combined results from the English and BSL grammaticality judgement tasks (EGJT and BSLGJT) to create a single, modality-independent measure of language proficiency in the deaf group. Accuracy scores in the EGJT (%correct; mean=83.51, SD=11.4, N=25) and BSLGJT (mean=77.88, SD=13.1, N=21) were transformed into z-scores separately for each test. For each participant, the EGJT and BSLGJT z-scores were then compared, and the higher one was chosen for a combined modality-independent language proficiency score (Supplementary Figure 1).

### Multiple linear regression

Multiple linear regression analyses were conducted to investigate whether neural activity in the superior temporal cortex of deaf individuals can predict performance in the switching task. The data were analysed using a backward data entry method in JASP.^40^ The default stepping method criteria were used, where predictors with p < .05 are entered into the model and those with p > 0.1 are removed until all predictors fall within this criteria. SwitchCost_RT_ and SwitchCost_ACC_ were entered as dependent variables in separate analyses. Each regression analysis had three covariates: neural switch cost in the right hemisphere, neural switch cost in the left hemisphere, and language.

Neural switch cost (BOLD_switch_ – BOLD_stay_) was calculated in ROIs with significant differences between the switch and stay condition in the deaf group. The average neural activity in all stay trials (BOLD_stay_) was subtracted from the average activity in the first switch trials (BOLD_switch_), and then averaged across ROIs separately in the right and left hemisphere.

### Data Sharing

Link to data and analysis files: https://osf.io/uh2ap/

## Results

### Behavioural results

Deaf (N=25) and hearing (N=20) individuals were scanned while performing four executive function tasks: working memory, planning, switching, and inhibition (Figure 1). Behavioural results from all tasks are shown in Figure 3. To explore differences in performance between groups, we conducted 2×2 repeated-measures ANOVAs for each task, with either accuracy or reaction time (RT) as the dependent variable, between-subjects factor group (hearing, deaf), and within-subjects factor condition (HEF, LEF). Results show a significant main effect of condition for both accuracy and RT in all tasks, confirming that the HEF condition was more difficult and demanding than the LEF condition (Supplementary Table 5).

**Figure 3.**
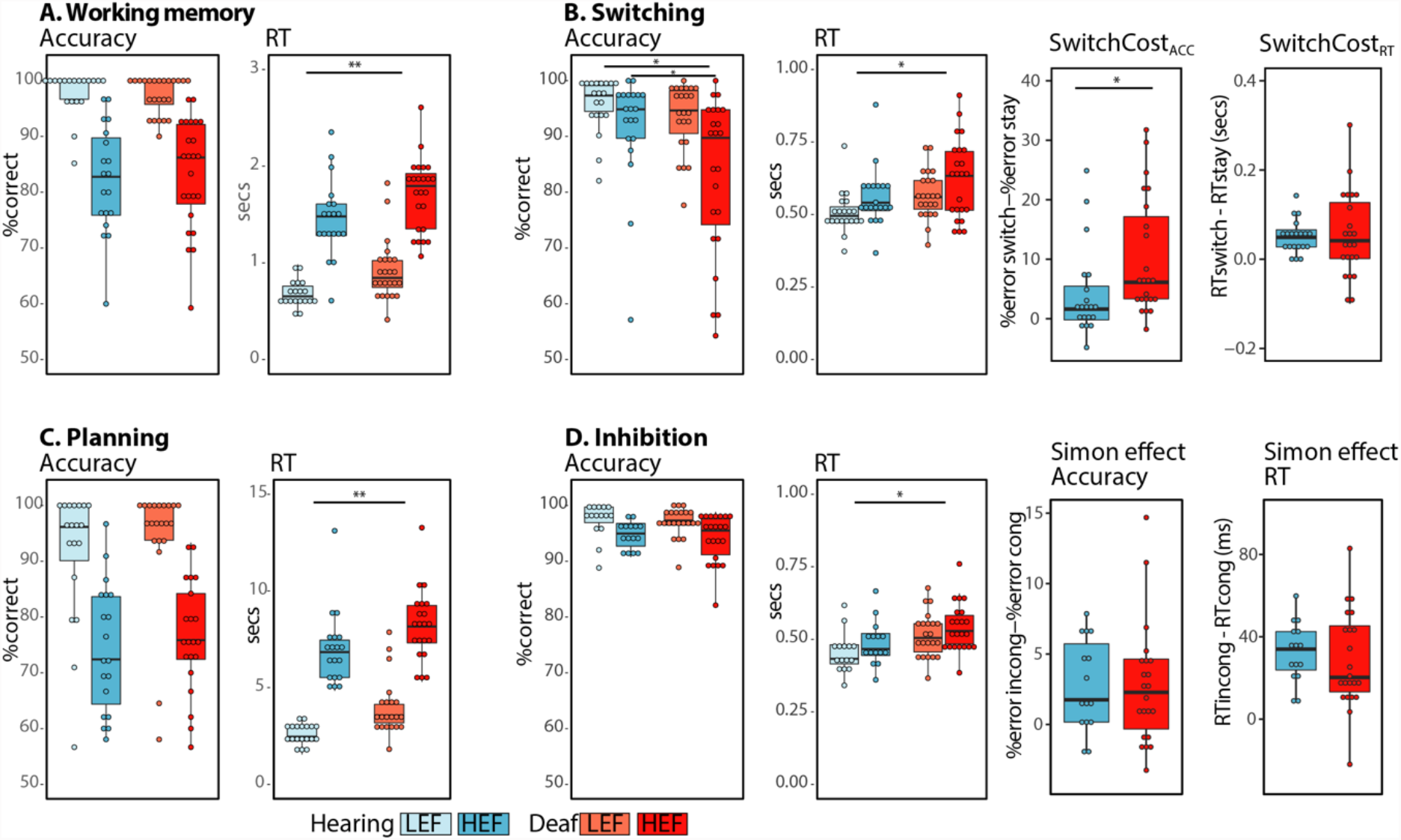
Behavioural performance. The figure shows average accuracy (%correct) and reaction time (seconds) for each task and condition in the hearing and the deaf groups. It also shows the average switch costs and Simon effects for both accuracy and reaction time in each group. The SwitchCost_ACC_ and Simon effect are calculated and plotted using %error instead of %correct, so that larger values indicate an increase in cost. Only the first trials of the switch blocks were included in the HEF condition. The bold lines in the box plots indicate the median. The lower and upper hinges correspond to the first and third quartiles. Statistically significant (p<0.05) differences between conditions are not shown in the figure, but were found for all tasks in both groups (Suppl. Table 5). **p<0.01; *p<0.05.

The group of deaf individuals had significantly slower RTs in all tasks (Supplementary Table 5).

Switching was the only task where there was a significant main effect of group on accuracy (F_1,41_=4.32, p=0.04), as well as a condition × group interaction (F_1,41_=4.98, p=0.03). A post-hoc t-test revealed that the deaf group was significantly less accurate in the switch condition (t_41_=-2.22, p=0.03). The difference in SwitchCost_ACC_ (%errors_switch_–%errors_stay_) reflects the significant interaction, with the deaf group (mean=10.24; SD=9.89) having a larger SwitchCost_ACC_ than the hearing group (mean=4.18; SD=7.53; t_41_=2.23, p=0.03; Figure 3B).

### Task switching activates temporal auditory areas in deaf individuals and this activation predicts behavioural performance

fMRI results show that all executive function tasks activated typical frontoparietal regions in both groups of participants (Supplementary Figure 2). To investigate crossmodal plasticity and executive processing in the deaf auditory cortex, we conducted a region of interest (ROI) analysis on superior temporal auditory ROIs. These included: Heschl’s gyrus (HG), the planum temporale (PT), and the posterior superior temporal cortex (pSTC) (Figure 2).

Of the four tasks that we tested, only in the switching task we found both a significant main effect of group (F_1,41_=15.48, p<0.001) and a significant interaction between group x condition (F_1,41_=4.75, p=0.03) (Table 1). The interaction was driven by a significant difference between conditions in the deaf group, but not in the hearing group (deaf_HEFvLEF_: t_22_=4.06, p=0.0005; hearing_HEFvLEF_: t_19_=0.26, p=0.27). To test whether differences between conditions were significant between the switch and stay condition in all ROIs, we conducted post-hoc t-tests in each ROI and group. This accounted for a total of 12 separate t-tests, and to correct for multiple comparisons, we only considered significant those results with p < .004 (p < .05/12 = .004; corrected p < .05). We found significant differences between the switch and stay condition in all the left hemisphere ROIs and in the right PT and right pSTC in the deaf group (Figure 4; Supp. Table 6).

**Table 1.** Group main effects and Group interactions for all tasks in the ROIs analysis.

|  | **Working Memory** |  | **Switching** |  | **Planning** |  | **Inhibition** |  |
| --- | --- | --- | --- | --- | --- | --- | --- | --- |
|  | *F (df)* | *p* | *F (df)* | *p* | *F (df)* | *p* | *F (df)* | *p* |
| **Group** | 0.038<br>(1,41) | 0.85 | **15.48**<br>(1,41) | **<0.001** | **5.85**<br>(1,38) | **0.02** | 0.03<br>(1,35) | 0.87 |
| **Condition × Group** | **6.40**<br>(1,41) | **0.01** | **4.75**<br>(1,41) | **0.03** | 0.56<br>(1,38) | 0.46 | 0.18<br>(1,35) | 0.67 |
| **ROI × Group** | 1.18<br>(1.67, 68.42) | 0.30 | **3.42**<br>(1.93, 79.05) | **0.04** | 0.73<br>(1.70, 64.55) | 0.46 | **3.92**<br>(1.89, 66.05) | **0.03** |
| **Hemisphere × Group** | 0.01<br>(1, 41) | 0.93 | 0.009<br>(1, 41) | 0.92 | 0.46<br>(1, 38) | 0.50 | 0.30<br>(1, 35) | 0.59 |

Full results for each ANOVA can be found in OSF: https://osf.io/dt827/

**Figure 4.**
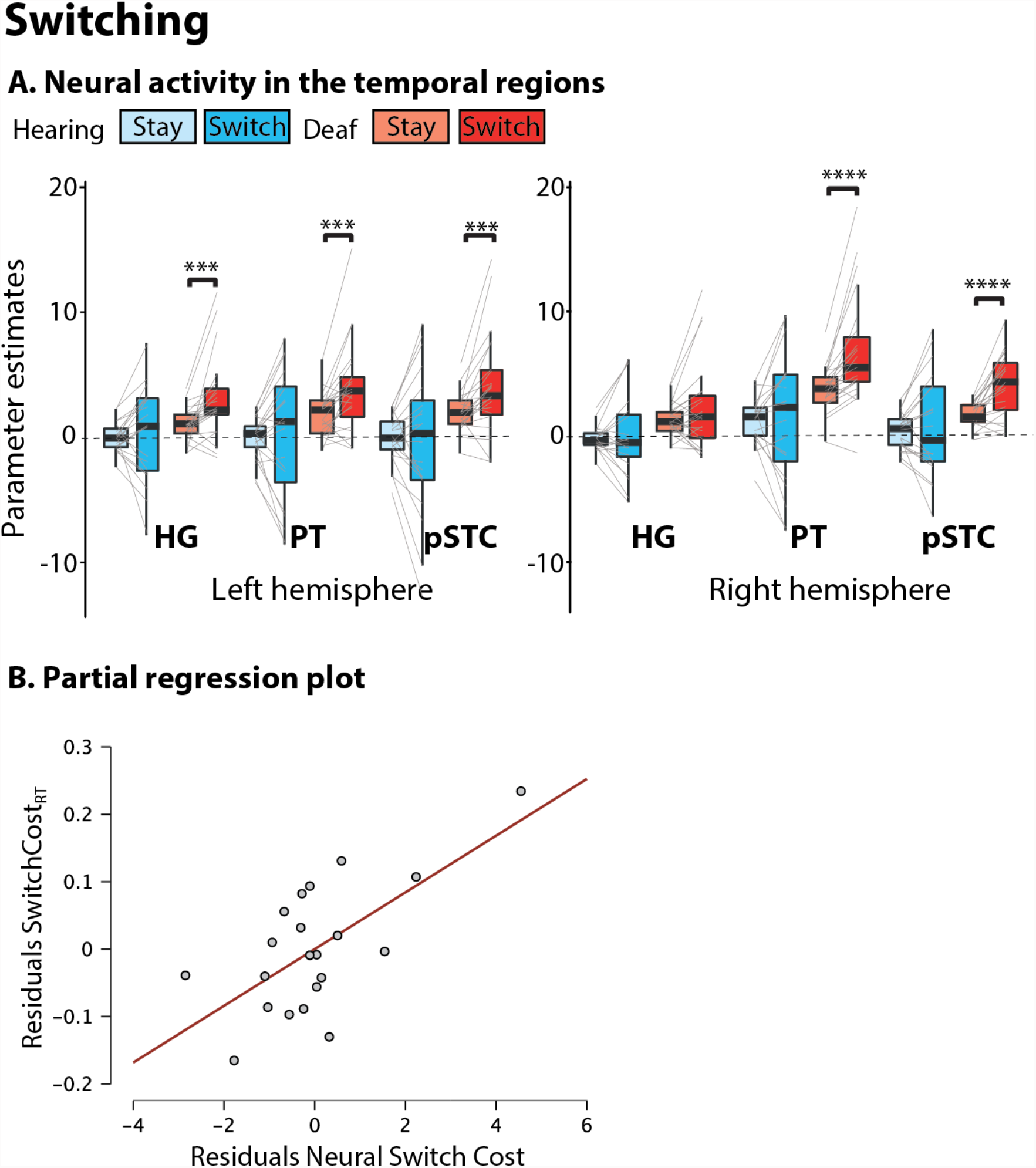
Switching task analysis. **A**. Neural activity in temporal ROIs. ***p<0.005; ****p<0.001. **B**. Partial correlation plot between SwitchCost_RT_ and neural switch cost in right temporal ROIs. Partial correlation from a multiple linear model with SwitchCost_RT_ as dependent variable and the following covariates: right hemisphere neural switch cost, left hemisphere neural switch cost, and language.

To investigate the behavioural relevance of the observed plasticity, we evaluated whether neural activity in the superior temporal cortex of deaf individuals can predict performance during the switching task. We conducted two separate multiple linear regression analyses, one with SwitchCost_RT_ and one with SwitchCost_ACC_ as dependent variables (Table 2).

**Table 2.** Multiple Linear Regression predicting behavioural performance in the switching task.

| SwitchCost<sub>RT</sub> |  |  |  |  |  |  |
| --- | --- | --- | --- | --- | --- | --- |
| Model Summary |  |  |  |  |  |  |
| Model |  | R<sup>2</sup> | Adjusted R<sup>2</sup> | F | p |  |
| 1 |  | 0.457 | 0.362 | 4.78 | 0.014 |  |
| 2 |  | 0.406 | 0.340 | 6.15 | 0.009 |  |
| Coefficients |  |  |  |  |  |  |
| Model |  | Unstandardized | Standard Error | Standardized | t | p |
| 1 | (Intercept) | 0.025 | 0.040 |  | 0.635 | 0.534 |
|  | Language Score | -0.061 | 0.048 | -0.232 | -1.270 | 0.221 |
|  | LH Neural Switch Cost | -0.024 | 0.010 | -0.693 | -2.350 | 0.031 |
|  | RH Neural Switch Cost | 0.042 | 0.012 | 1.053 | 3.605 | 0.002 |
| 2 | (Intercept) | -0.013 | 0.026 |  | -0.511 | 0.616 |
|  | LH Neural Switch Cost | -0.021 | 0.010 | -0.613 | -2.093 | 0.051 |
|  | RH Neural Switch Cost | 0.040 | 0.012 | 0.992 | 3.387 | 0.003 |

| SwitchCost<sub>ACC</sub> |  |  |  |  |  |  |
| --- | --- | --- | --- | --- | --- | --- |
| Model Summary |  |  |  |  |  |  |
| Model |  | R<sup>2</sup> | Adjusted R<sup>2</sup> | F | p |  |
| 1 |  | 0.285 | 0.159 | 2.26 | 0.119 |  |
| 2 |  | 0.283 | 0.203 | 3.55 | 0.050 |  |
| 3 |  | 0.207 | 0.165 | 4.96 | 0.038 |  |
| Coefficients |  |  |  |  |  |  |
| Model |  | Unstandardized | Standard Error | Standardized | t | p |
| 1 | (Intercept) | 15.844 | 4.823 |  | 3.285 | 0.004 |
|  | Language Score | -12.850 | 5.834 | -0.462 | -2.203 | 0.042 |
|  | LH Neural Switch Cost | -0.282 | 1.249 | -0.076 | -0.225 | 0.824 |
|  | RH Neural Switch Cost | 1.410 | 1.411 | 0.335 | 1.000 | 0.331 |
| 2 | (Intercept) | 15.779 | 4.686 |  | 3.368 | 0.003 |
|  | Language Score | -12.570 | 5.547 | -0.452 | -2.266 | 0.036 |
|  | RH Neural Switch Cost | 1.159 | 0.840 | 0.275 | 1.380 | 0.185 |
| 3 | (Intercept) | 18.900 | 4.200 |  | 4.500 | < .001 |
|  | Language Score | -12.644 | 5.677 | -0.455 | -2.227 | 0.038 |

LH: left hemisphere; RH: right hemisphere.

In the ROI analysis, there was a significant difference in activity between the switch and stay conditions in the deaf group in seven ROIs. To reduce the number of covariates and the dimensions of the models, we calculated the neural switch cost (BOLD_switch_ – BOLD_stay_) for each ROI, and then averaged across ROIs in the right and left hemisphere separately. We included language as a covariate in our model, as language proficiency has been shown to modulate performance in EF tasks in deaf individuals.^44–47^

Multiple linear regression using backward data entry shows that neural activity in temporal ROIs can significantly predict SwitchCost_RT_ in the deaf group, with the most significant model explaining 40.6% of the variance (F(2,18)=6.15, p=.009 R2= 0.406, adjusted R2=0.34; Table 2, Top panel). There was a positive association between neural switch cost in right hemisphere temporal areas and SwitchCost_RT_ (B= 0.04, SE= 0.01, *β*=0.99; p=.003). This means that for every unit increase in neural switch cost in right temporal areas, there is an increase of 40ms in SwitchCost_RT_. In standardised terms, as neural switch cost increases by 1 standard deviation, SwitchCost_RT_ increases by 0.99 SDs. On the other hand, there was a negative association between the left hemisphere neural and SwitchCost_RT_, and while this was significant at p=.031 in the full model, the significance was only p=.051 in the best model (Table 2). There was no significant association between language and SwitchCost_RT_ (B= -0.06, SE= 0.048, *β*=-0.23; p=.22).

When evaluating whether neural switch cost could also predict SwitchCost_ACC_, we found no significant association between these variables (Table 2, Bottom panel). Instead, the multiple linear regression shows that only language can significantly predict SwitchCost_ACC_, with the most significant model explaining 20.7% of the variance at p=.038 (F(1,19)=4.96, p=.038, R^2^=0.207, adjusted R^2^=.165).

### Recruitment of auditory areas in deaf individuals is not ubiquitous across EF tasks

Results from the working memory, planning and inhibition tasks are shown in Figure 5. In the working memory task, there was a significant condition × group interaction (Table 1, F_1,41_=6.41, p=0.01), but differences between conditions within each group were not significant (hearing_HEFvLEF_: t_18_=1.74, p=0.10; deaf_HEFvLEF_: t_23_=1.81, p=0.08). In the planning task, there was a significant main effect of group (F_1,38_=5.85, p=0.02), but this was driven by significant deactivations in the hearing group (t_18_=-4.47, p<0.001), with no significant difference in activity from baseline in the deaf group (t_20_=-1.31, p=0.21). In the Inhibition task, there was a significant interaction between ROI and Group (F_1.89,66.05_=3.92, p=0.03). However, there were no significant differences between groups in any ROI (https://osf.io/9fuec). Instead, the ROI x group interaction was driven by a main effect of ROI in the deaf group (higher activations for PT and pSTC than HG, https://osf.io/2z35e/), which was not present in the hearing group (https://osf.io/gmy6v/).

**Figure 5.**
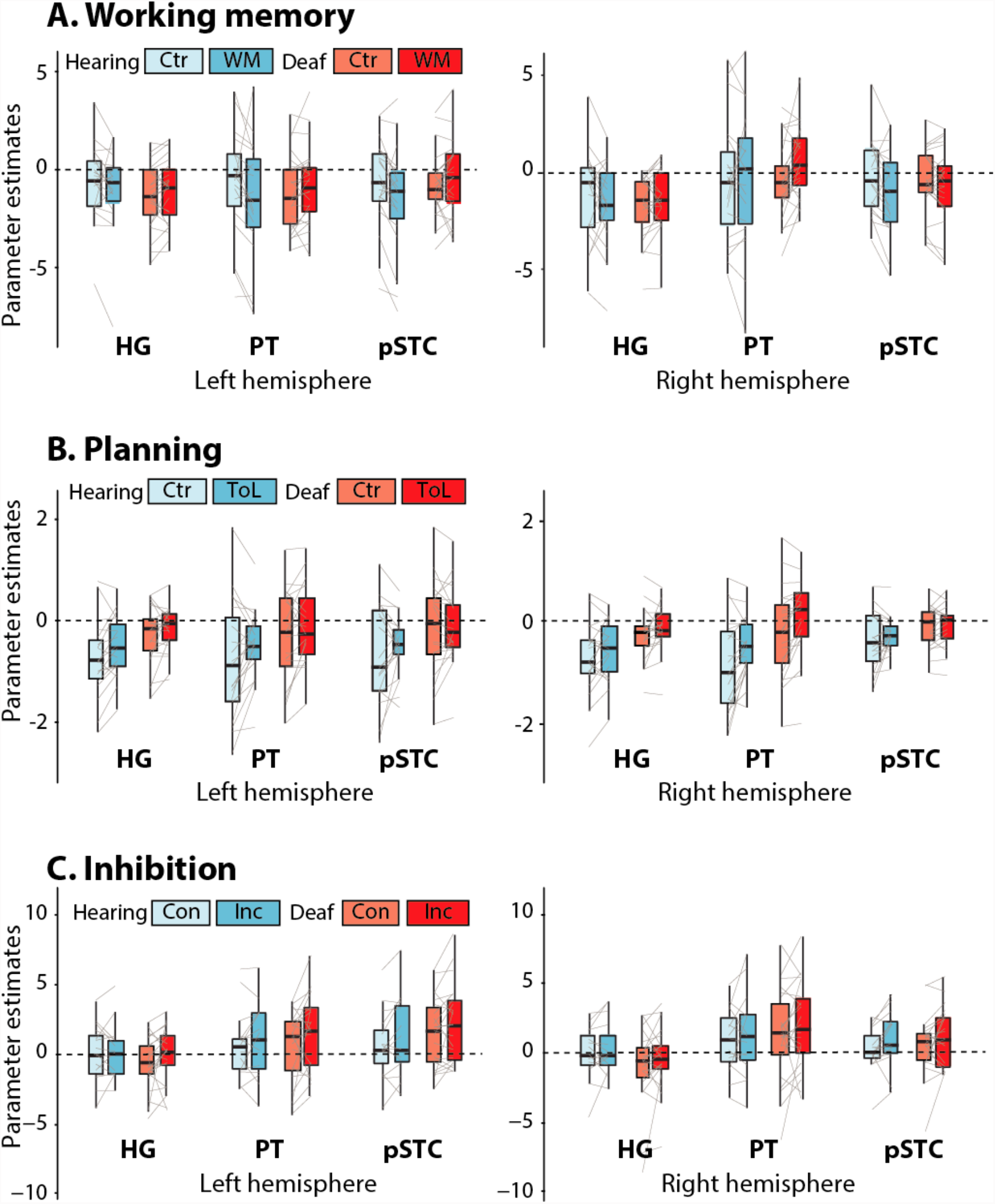
ROI results from the working memory, planning and inhibition tasks. Ctr=control, WM=working memory, ToL=Tower of London, Con=congruent, Inc=incongruent. HG=Heschl’s gyrus, PT=planum temporale, pSTC=posterior superior temporal cortex.

## Discussion

We investigated how early sensory experience impacts the organisation of executive processing in the brain. We found that, in deaf individuals, primary and secondary auditory areas are recruited during a visual switching task. These results suggest that the sensory or cognitive specialisation of cortical regions in the adult brain can be influenced by developmental sensory experience. It is possible that an early absence of auditory inputs results in a shift of functions in regions typically involved in auditory processing, with these regions then adopting a role in specific components of executive processing. Neural activity in temporal regions during the switching task predicted performance in deaf individuals, highlighting the behavioural relevance of this functional shift.

Our design allowed us to thoroughly examine the role of auditory regions in different executive function tasks and determine whether these regions are involved in cognitive control. Previous studies have suggested an involvement of auditory cortex during higher-order cognitive tasks in deaf individuals^3,4^, but given the focus on a single task, with an experimental and control condition, they cannot inform whether plasticity effects are specific to the demands of the task. Our design included four different visuospatial EF tasks, all with an experimental (HEF) and control (LEF) condition, probing a variety of executive processes. We found that the HEF condition in all tasks recruited frontoparietal areas typically involved in executive functioning and cognitive control. However, only switching resulted in significant activations in temporal auditory regions in the deaf group. This finding demonstrates that the deaf auditory cortex serves a specific subcomponent of executive functioning during switching, and not a shared computation across tasks, such as cognitive control. This was not only found in higher-order auditory areas, but also in the left Heschl’s Gyrus, showing that a functional shift towards cognition can indeed occur in primary sensory regions.^48^

Task switching requires cognitive flexibility and shifting between different sets of rules.^49,50^ Shifting is considered one of the core components of executive control. It is defined as the ability to flexibly shift “back and forth between multiple tasks, operations, or mental sets”.^51^ While all four tasks were cognitively demanding, only the switching task involved both frequent and rapid shifts between arbitrary stimulus-response mappings as well as speeded manual responses. This specific combination of fast task shifting and rapid response selection and execution made the switching task temporally demanding. We suggest that such high temporal task demands may particularly suit the “auditory” cortex, given the critical importance of fine-grained temporal analysis for the cortical processing of speech and other complex sounds.^48^ Shifting is also an important component of working memory tasks previously shown to recruit posterior superior temporal regions in deaf individuals (e.g. 2-back working memory, visuospatial delayed recognition^3,4^). In the present study, the working memory task did not significantly activate any temporal ROI. The working memory task used in this study requires updating of information and incremental storage, but no shifting between targets or internal representations of stimuli, as required in an n-back task. Together, these results suggest that previous working memory effects in superior temporal regions are not necessarily linked to storage, updating or control, but are more likely linked to shifting between tasks or mental states.

A possible physiological mechanism supporting this change of function in the auditory cortex, specifically in the right hemisphere, can be provided through anatomical proximity to the middle temporal lobe or to the parietal lobe, specifically the temporoparietal junction (TPJ).^6,52^ Right TPJ is a multisensory associative region involved in reorientation of attention to task-relevant information, such as contextual cues or target stimuli.^53,54^ Regions of the right middle temporal gyrus have also been shown to be involved in task switching^55^ and to encode task-set representations.^56^ This is particularly relevant because right hemisphere auditory regions were more strongly linked to behavioural outcomes in the deaf group. The anatomical location and the functional role of right TPJ and the middle temporal gyrus suggest that, in the absence of auditory inputs throughout development, the computations performed by these regions could be extended to adjacent auditory cortices.^6,52,57^

Another possibility is that the recruitment of “auditory” temporal regions for switching observed in deaf adults reflects vestigial functional organisation present in early stages of development. Research on hearing children has found activations in bilateral occipital and superior temporal cortices during task switching^58^, with a similar anatomical distribution to the one we find here. Our findings in deaf individuals suggest that executive processing in temporal cortices could be “displaced” by persistent auditory inputs which, as the individual develops, may require more refined processing or demanding computations. Thus, an alternative view is that regions considered to be “sensory” have mixed functions in infants and become more specialised in adults. These regions could follow different developmental pathways influenced by environmental sensory experience. As such, the temporal regions of hearing individuals will become progressively more specialised for sound processing, whereas, in deaf individuals, they will become more specialised for subcomponents of executive processing.

The direct relationship between behavioural outcomes and activity in reorganised cortical areas is robust evidence of the functional importance of the observed crossmodal plasticity. We found that neural activity, specifically in the right temporal ROIs, predicted reaction times in the switching task in the deaf group. Specifically, higher neural switch cost was linked to higher RT switch cost (SwitchCost_RT_), which suggests effortful processing, as previously described in other cognitive tasks with different levels of complexity.^59,60^ It is important to highlight that there were no differences in SwitchCost_RT_ between the groups, showing that the potential reliance on different neural substrates to solve the switching task does not translate into differences in performance. In fact, significant interactions between group and condition for the switching task were only found in accuracy (SwitchCost_ACC_), which in our analysis was not predicted by neural activity, but rather, by language proficiency. Executive performance has been previously associated with language proficiency in deaf children.^46,47,61–63^ While in our study language z-scores predict only 20.7% of the variance in SwitchCost_ACC_ and the model was only significant at p <0.05, our findings suggest that language development can have long-lasting effects on executive processing throughout the lifespan. Different theories propose that language can provide the necessary framework for higher-order (if-if-then) rules to develop and be used in a dynamic task in the most efficient way.^64,65^ These hierarchical “if-then” rules could be implemented, in an automatic way, to solve the arbitrary link between stimulus and response during switching. Although participants are not required to use linguistic strategies during switching, we speculate that those who have benefited from the efficiency associated with developing such frameworks can invest less cognitive resources into solving this task. While the role of language in executive processing needs further investigation, it is important to consider that the timely development of a first language may boost the overall efficiency of a cognitive task, in this case switching, regardless of whether the task itself allows implementation of purely linguistic mechanisms.

In conclusion, we show that executive processing in the adult brain is influenced by early sensory experience. Our results suggest that the absence of auditory inputs could “free” superior temporal regions to take on functions other than sensory processing. This could be either by preserving a function these areas performed early in childhood or by taking on new functions driven by influences from top-down projections from frontoparietal areas or adjacent temporal and parietal regions.

## Acknowledgments

The authors would like to specially thank all the deaf and hearing participants who took part in this study. This work was funded by a grant from the Biotechnology and Biological Sciences Research Council (BBSRC; BB/P019994). VV is funded by a scholarship from the University of East Anglia.

## Supplementary Figures

**Supplementary Figure 1.**
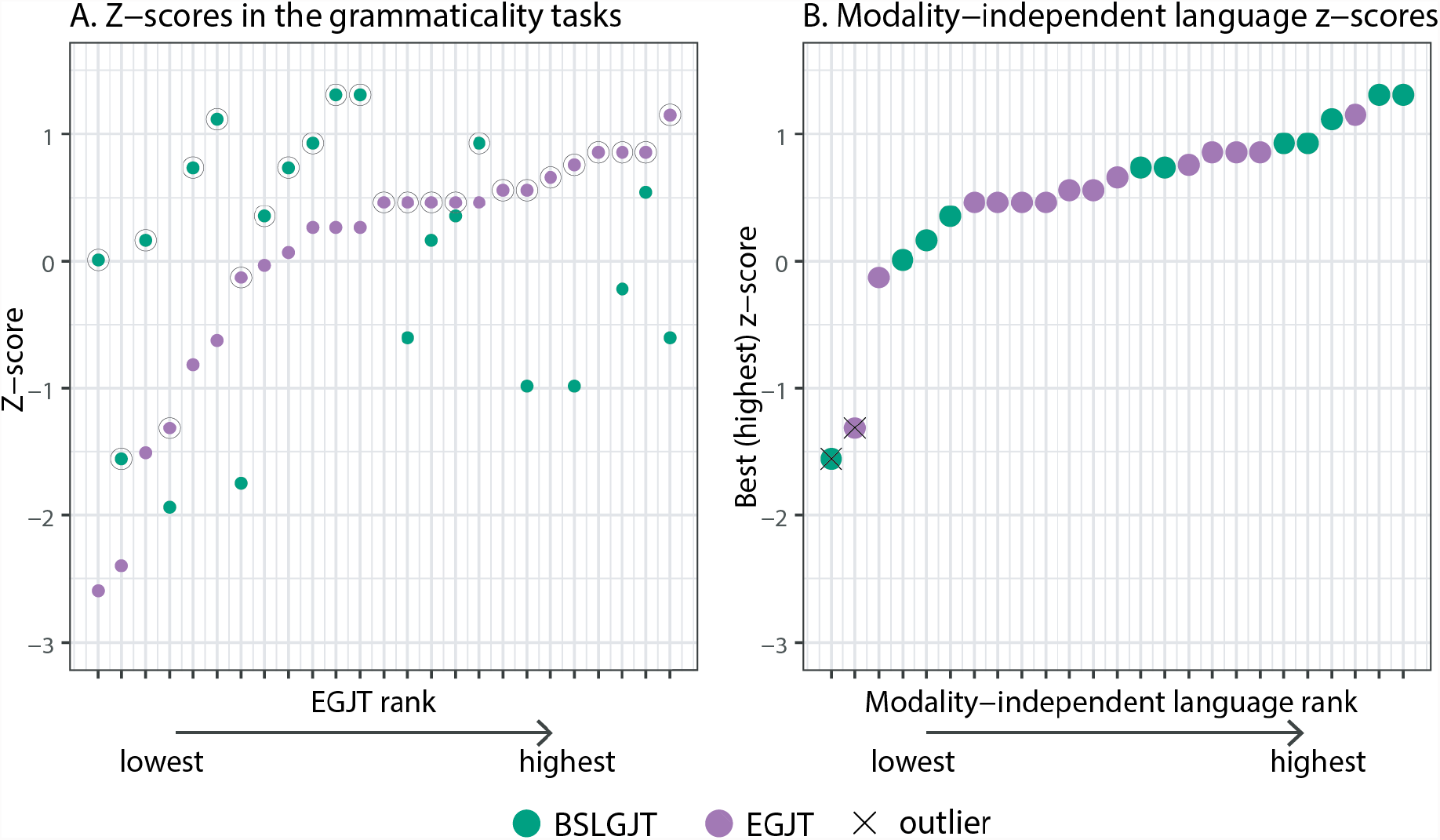
Language proficiency in the deaf group. **A** Language z-scores in the English grammaticality judgement task (EGJT) and BSL grammaticality judgement task (BSLGJT), with participants sorted on the x-axis by their EGJT rank. Black circles indicate the z-score chosen for the combined modality-independent language score.

**Supplementary Figure 2.**
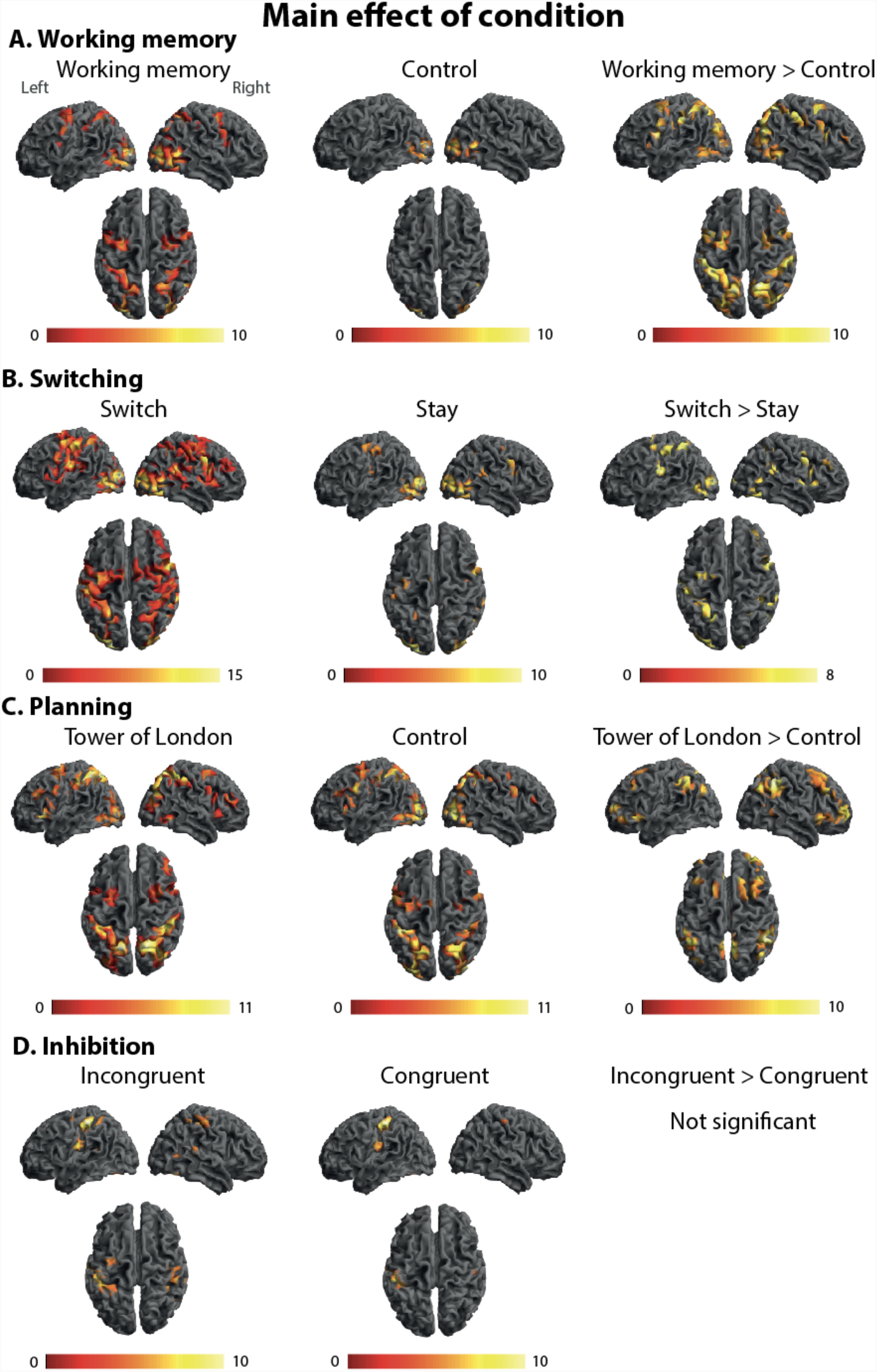
Whole-brain main effects of condition in each EF task. Activations for each EF task and condition averaged across groups. There were significantly stronger activations in the HEF condition in the working memory, planning, and switching tasks. These included commonly found activations in frontoparietal areas, such as dorsolateral prefrontal cortex (DLPFC), frontal eye fields (FEF), pre-supplementary motor area (pre-SMA), and intraparietal sulcus (IPS). In the inhibition task, the HEF incongruent condition resulted in stronger activation in IPS and left FEF, but there were no significant differences between conditions. Contrasts calculated across both groups. All contrasts are displayed at p<0.05 (FWE-corrected). Colour bars represent z-scores. Coordinates of peak activations for the contrast [HEF > LEF] can be found here: https://osf.io/87vur/

## Supplementary Tables

**Supplementary Table 1.** Demographics and pre-screening tests

|  | Age |  | Gender | WASI |  | Corsi |  |
| --- | --- | --- | --- | --- | --- | --- | --- |
|  | Mean (range) | SD |  | Mean | SD | Mean | SD |
| **Full sample**<br>hearing N=20 | 37.50 (18-66) | 16.85 | 15f/5m | 57.47 | 8.02 | 5.4 | 1.1 |
| **Full sample**<br>deaf N=25 | 41.68 (19-66) | 14.38 | 16f/9m | 59.68 | 8.51 | 5.3 | 0.78 |
| **WM**<br>hearing N=19 | 38.47 (18-66) | 16.72 | 14f/5m | 57.83 | 8.1 | 5.47 | 1.08 |
| **WM**<br>deaf N=24 | 41.38 (19-66) | 14.6 | 15f/9m | 59.96 | 8.58 | 5.38 | 0.70 |
| **Planning**<br>hearing N=19 | 36.95 (18-66) | 17.13 | 14f/5m | 57.56 | 8.25 | 5.44 | 1.11 |
| **Planning**<br>deaf N=21 | 40.81 (19-63) | 13.65 | 13f/8m | 59.67 | 9.19 | 5.26 | 0.82 |
| **Switching**<br>hearing N=20 | 37.50 (18-66) | 16.85 | 15f/5m | 57.47 | 8.02 | 5.4 | 1.1 |
| **Switching**<br>deaf N=23 | 40.3 (19-63) | 13.93 | 14f/9m | 59.87 | 8.76 | 5.39 | 0.71 |
| **Inhibition**<br>hearing N=15 | 40.33 (18-66) | 17.09 | 12f/3m | 57.57 | 9.05 | 5.43 | 1.21 |
| **Inhibition**<br>deaf N=22 | 40.59 (19-66) | 14.87 | 14f/8m | 60.05 | 8.84 | 5.43 | 0.70 |

**Supplementary Table 2.** Between-group comparisons on demographics and pre-screening tests

|  | **N** | **N** | **Age** |  |  | **Gender** |  | **WASI** |  |  | **Corsi** |  |  |
| --- | --- | --- | --- | --- | --- | --- | --- | --- | --- | --- | --- | --- | --- |
| | hearing | deaf | *df* | *t* | *p* | $\chi^2$ | *p* | *df* | *t* | *p* | *df* | *t* | *p* |
| **WM** | 19 | 24 | 1,41 | 0.6 | 0.55 | 0.6 | 0.44 | 1,40 | 0.81 | 0.42 | 1,40 | -0.36 | 0.72 |
| **Planning** | 19 | 21 | 1,38 | 0.79 | 0.43 | 0.63 | 0.43 | 1,37 | 0.75 | 0.46 | 1,37 | -0.59 | 0.56 |
| **Switching** | 20 | 23 | 1,41 | 0.6 | 0.55 | 0.97 | 0.32 | 1,40 | 0.92 | 0.37 | 1,40 | -0.01 | 0.99 |
| **Inhibition** | 15 | 22 | 1,35 | 0.05 | 0.96 | 1.14 | 0.29 | 1,34 | 0.81 | 0.42 | 1,34 | 0.01 | 0.99 |

See Supplementary Table 4 for exclusion criteria from individual tasks.

**Supplementary Table 3.**
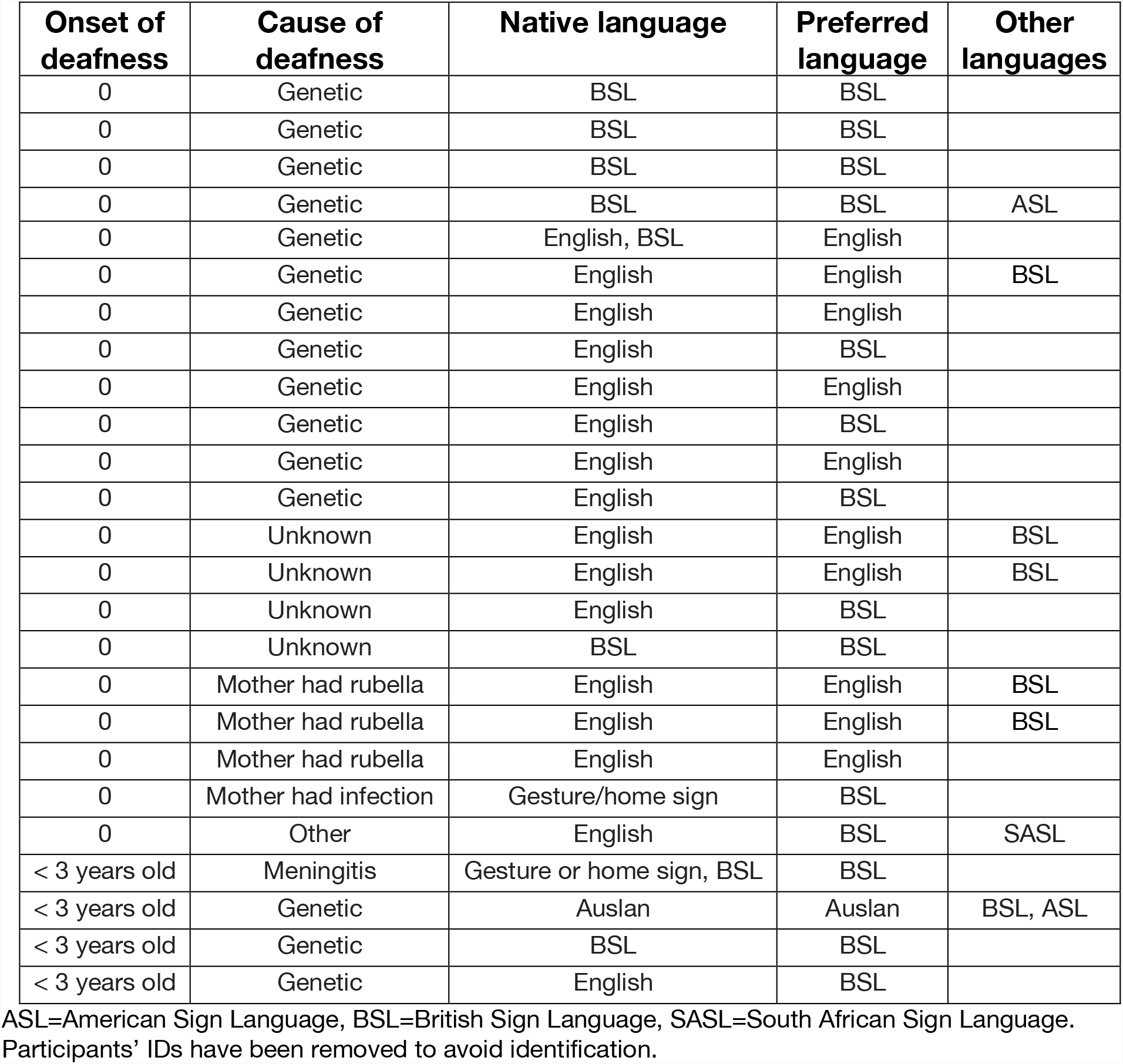
Cause of deafness and language background of the deaf participants

| **Onset of deafness** | **Cause of deafness** | **Native language** | **Preferred language** | **Other languages** |
| --- | --- | --- | --- | --- |
| 0 | Genetic | BSL | BSL |  |
| 0 | Genetic | BSL | BSL |  |
| 0 | Genetic | BSL | BSL |  |
| 0 | Genetic | BSL | BSL | ASL |
| 0 | Genetic | English, BSL | English |  |
| 0 | Genetic | English | English | BSL |
| 0 | Genetic | English | English |  |
| 0 | Genetic | English | BSL |  |
| 0 | Genetic | English | English |  |
| 0 | Genetic | English | BSL |  |
| 0 | Genetic | English | English |  |
| 0 | Genetic | English | BSL |  |
| 0 | Unknown | English | English | BSL |
| 0 | Unknown | English | English | BSL |
| 0 | Unknown | English | BSL |  |
| 0 | Unknown | BSL | BSL |  |
| 0 | Mother had rubella | English | English | BSL |
| 0 | Mother had rubella | English | English | BSL |
| 0 | Mother had rubella | English | English |  |
| 0 | Mother had infection | Gesture/home sign | BSL |  |
| 0 | Other | English | BSL | SASL |
| < 3 years old | Meningitis | Gesture or home sign, BSL | BSL |  |
| < 3 years old | Genetic | Auslan | Auslan | BSL, ASL |
| < 3 years old | Genetic | BSL | BSL |  |
| < 3 years old | Genetic | English | BSL |  |

ASL=American Sign Language, BSL=British Sign Language, SASL=South African Sign Language. Participants' IDs have been removed to avoid identification.

**Supplementary Table 4.**
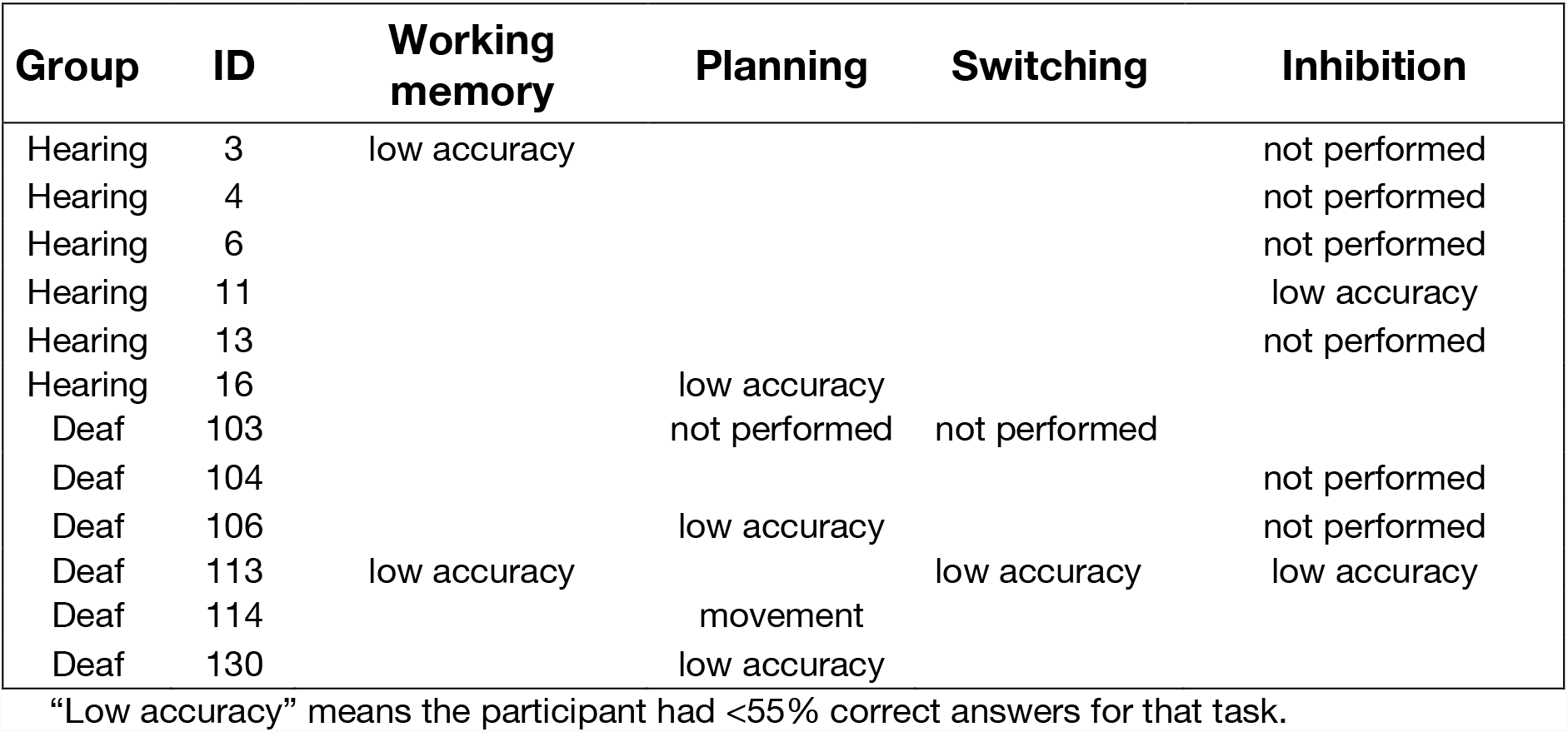
List of participants excluded from the analysis of each task

**Supplementary Table 5.** Results from repeated-measures ANOVAs on behavioural performance

|  | WM |  | Switching |  | Planning |  | Inhibition |  |
| --- | --- | --- | --- | --- | --- | --- | --- | --- |
|  | Accuracy |  |  |  |  |  |  |  |
|  | *F (df)* | *p* | *F (df)* | *p* | *F (df)* | *p* | *F (df)* | *p* |
| Condition | 91.59 (1,41) | <0.001 | 28.27 (1,41) | <0.001 | 46.88 (1,38) | <0.001 | 17.57 (1,35) | <0.001 |
| Group | 0.03 (1,41) | 0.86 | 4.32 (1,41) | 0.04 | 0.86 (1,38) | 0.36 | 0.24 (1,35) | 0.63 |
| Condition × Group | 0.3 (1,41) | 0.59 | 4.98 (1,41) | 0.03 | 0.01 (1,38) | 0.92 | 0.00 (1,35) | 0.98 |
|  | Reaction time |  |  |  |  |  |  |  |
|  | *F (df)* | *p* | *F (df)* | *p* | *F (df)* | *p* | *F (df)* | *p* |
| Condition | 199.22 (1,41) | <0.001 | 21.6 (1,41) | <0.001 | 211.64 (1,38) | <0.001 | 79.2 (1,35) | <0.001 |
| Group | 8.11 (1,41) | 0.007 | 4.5 (1,41) | 0.04 | 10.96 (1,38) | 0.002 | 4.91 (1,35) | 0.03 |
| Condition × Group | 0.0 (1,41) | 0.97 | 0.03 (1,41) | 0.87 | 0.06 (1,38) | 0.8 | 0.35 (1,35) | .56 |

Factors in the analysis were: condition (HEF, LEF) and group (hearing, deaf). Significant effects are shown in bold.

**Supplementary Table 6.** Paired samples t-tests (switch v stay) for each group and ROI in the switching task.

|  | Hearing group<br>*df= 19* |  |  |  | Deaf group<br>*df= 22* |  |  |  |
| --- | --- | --- | --- | --- | --- | --- | --- | --- |
|  | Left hemisphere |  | Right hemisphere |  | Left hemisphere |  | Right hemisphere |  |
|  | *t* | *p* | *t* | *p* | *t* | *p* | *t* | *p* |
| Heschl's gyrus | 0.41 | 0.68 | 0.35 | 0.73 | **3.38** | **0.003** | 1.58 | 0.13 |
| Planum temporale | 0.21 | 0.84 | 0.27 | 0.79 | **3.55** | **0.002** | **4.55** | **<0.001** |
| Posterior superior temporal cortex | 0.22 | 0.82 | 0.58 | 0.57 | **3.32** | **0.003** | **5.31** | **<0.001** |

Significant results are shown in bold. The following correction for multiple comparison was applied: results are considered significant when $p < 0.004$ ( $p < 0.05/12 = 0.004$ ; corrected $p < 0.05$ ).

## Notes

https://osf.io/uh2ap/

### Competing Interest Statement

The authors have declared no competing interest.

### Summary of Updates

The paper has been re-structured to focus solely on crossmodal plasticity in auditory areas. Results on the relationship between language experience and the reorganisation of cognitive networks of the brain will be reported in a separate manuscript. Errors in the behavioural analysis of the inhibition task have been corrected. This has not changed the conclusions of the paper.

